# Chemogenetics reveal an anterior cingulate-thalamic pathway for attending to task-relevant information

**DOI:** 10.1101/2020.06.22.164996

**Authors:** Emma J. Bubb, John P. Aggleton, Shane M. O’Mara, Andrew J.D. Nelson

## Abstract

In a changing environment, we need to decide when to select items that resemble previously rewarded stimuli and when it is best to switch to other stimulus types. Here, we used chemogenetic techniques to provide causal evidence that activity in the rodent anterior cingulate cortex and its efferents to the anterior thalamic nuclei modulate the ability to attend to reliable predictors of important outcomes. Rats were tested on an attentional set-shifting paradigm that first measures the ability to master serial discriminations involving a constant stimulus dimension that reliably predicts reinforcement (intradimensional-shift), followed by the ability to shift attention to a previously irrelevant class of stimuli when reinforcement contingencies change (extradimensional-shift). Chemogenetic silencing of the anterior cingulate cortex (Experiment 1) as well as selective inactivation of anterior cingulate efferents to the anterior thalamic nuclei (Experiment 2) impaired intradimensional learning but, facilitated two sets of extradimensional-shifts. This pattern of results signals the loss of a cortico-thalamic system for cognitive control that preferentially processes stimuli resembling those previously associated with reward. Previous studies highlight a separate prefrontal system that promotes switching to hitherto inconsistent predictors of reward when contingencies change. Competition between these two systems regulates cognitive flexibility and choice.

## Introduction

In a dynamic world, the ability to engage in adaptive behaviors is critical to an organism’s survival. This includes deciding when to select items that resemble consistently rewarded stimuli and when to switch to previously irrelevant stimulus types. The ability to disengage from previously rewarded response strategies depends on the integrity of prefrontal cortex. Consequently, prefrontal damage in humans, marmosets, and rats causes response perseveration and a failure to switch when contingencies change (Milner, 1963; Dias et al., 1996a, 1996b; Birrell and Brown, 2000; Ng et al., 2007; Nyhus and Barceló, 2009). Further research with rats highlights how interactions between prefrontal cortex and subcortical sites support this form of behavioral flexibility (Block et al., 2007; Baker and Ragozzino, 2014). However, until recently, there has been little progress in identifying the neural circuits in rodents that support the opposing attentional mechanism, i.e. the preferential processing of stimuli resembling those previously associated with important outcomes.

The attentional set-shifting paradigm captures both potentially conflicting cognitive processes. In this task, animals show accelerated learning over successive discriminations by attending to a common stimulus dimension (intradimensional-set) (Mackintosh, 1975). This learnt bias to a specific category (attentional-set) is further revealed when the stimulus being rewarded switches to a qualitatively different category. Now, additional trials, the ‘shift-cost’, are required to solve this extradimensional-shift (Birrell and Brown, 2000; Chase et al., 2012). Lesions in the rodent anterior thalamic nuclei (ATN) disrupt animals’ ability to form an attentional-set, as revealed by impaired intradimensional-set performance, but paradoxically, when required to solve discriminations involving hitherto irrelevant stimulus dimensions, lesion animals not only outperform controls, but display a shift-benefit (Wright et al., 2015).

The implication is that the ATN are vital for attending to reliable predictors of reinforcement, driving attentional-set formation at the expense of extradimensional-shifts. This bias is then lost following ATN lesions, resulting in heightened attention to inconsistent predictors of reward. This interpretation is supported by both functional imaging and clinical data in humans (de Bourbon-Teles et al., 2014; Leszczyński and Staudigl, 2016). The resulting dissociation between the effects of prefrontal lesions (impaired extradimensional-shift) and ATN lesions (facilitated extradimensional-shift) points to a distinct role for the ATN within cortico-thalamic circuits supporting attentional processes. A key question remains, therefore, with which cortical sites might the ATN act to support these processes?

A potential partner is the anterior cingulate cortex (ACC). The ATN are densely interconnected with the ACC (Shibata, 1993; Shibata and Naito, 2005; Wright et al., 2013) and there is evidence that the ACC contributes to attentional processing (Ng et al., 2007; Koike et al., 2016). For example, rats with ACC lesions oversample never-reinforced stimuli and appear more distracted by irrelevant information (Ragozzino and Rozman, 2007; Newman and McGaughy, 2011). To test this potential partnership, we first virally expressed the inhibitory hM4Di DREADD receptor in dorsal ACC. Rats then received behavioral assays explicitly designed to contrast attentional set-formation and extradimensional set-shifting (Chase et al., 2012; Lindgren et al., 2013). For these assays, the rats first received a series of two-choice discriminations based on either odors or digging media, and underwent a series of four intradimensional-shifts, during which one dimension (e.g. odor) is consistently rewarded, while the other dimension (e.g. media) remains irrelevant. Next, rats experienced an extradimensional-shift in which the previously irrelevant dimension is now rewarded. Subsequently, the rats performed a second extradimensional-shift task in which spatial position became, for the first time, relevant (Wright et al., 2015). Last, we interrogated the effects of DREADD activation by measuring c-*fos* expression in the ACC and relevant efferent targets.

Experiment 2 tested the specific hypothesis that interactions between the ACC and ATN support these attentional processes. Taking advantage of the anterograde transport of an adeno-associated virus expressing the inhibitory hM4Di DREADD receptor, coupled with localized infusions of the ligand within the ATN, we examined the effects of chemogentically silencing ACC efferents to the ATN on the same attentional set-shifting tasks.

## Materials and Methods

### Subjects

All experiments involved experimentally naïve, male Lister Hooded rats (Envigo, Bicester, UK). The rats were housed in pairs in a temperature-controlled room. At the time of surgery, the rats in Experiment 1 (n = 22) weighed between 290 and 331g, those in Experiment 2 (n = 18) weighed between 296 and 328g. Lighting was kept on a 12-hour light/dark cycle, light from 0800 to 2000. During behavioral testing all animals were food restricted to maintain at least 85% of their free-feeding body weight, while water was available *ad libitum*. All experiments were carried out in accordance with UK Animals (Scientific Procedures) Act, 1986 and EU directive (2010/63/EU) as well as local ethical approval from Cardiff University.

### Surgical Procedures

All rats were anesthetized with isoflurane (4% induction, 2% thereafter). Next, each rat was placed in a stereotaxic frame (David Kopf Instruments, Tujunga, CA), with the skull flat (Experiment 1) or with the incisor bar set at +5.0 to the horizontal plane (Experiment 2). For analgesic purposes, Lidocaine was administered topically to the scalp (0.1ml of 20mg/ml solution; B. Braun, Melsungen, Germany) and meloxicam was given subcutaneously (0.06ml of 5mg/ml solution, Boehringer Ingelheim Ltd, Berkshire, UK). A craniotomy was then made directly above the target region and the dura cut to expose the cortex.

In both experiments, the experimental group received injections of an adeno associated virus expressing the inhibitory hM4Di DREADD receptor into the anterior cingulate cortex while control animals received injections of the same virus not expressing the DREADD receptor. Injections were made via a 10μl Hamilton syringe (Bonaduz, Switzerland) attached to a moveable arm mounted to the stereotaxic frame. The injections were controlled by a microprocessor (World Precision Instruments, Hitchin, UK) set to a flow rate of 0.1 μl/min, and the needle left *in situ* for a further 5 minutes to allow for diffusion of the bolus.

In Experiment 1, the experimental group (n = 12) received injections of AAV5-CaMKIIa-hM4Di-mCherry (titer 4.4×10^12 GC/ml; Addgene, MA, USA;) and the control group (n = 10) received injections of AAV5-CaMKIIa-EGFP (titer 4.3×10^12 GC/ml; Addgene, MA, USA). The injection coordinates and volumes for the three injections made into the anterior cingulate cortex were as follows: 1) 0.35μl at AP +1.9, ML +/-0.8, DV −1.2; 2) 0.7μl at AP +1.0, ML +/- 0.8, DV −1.6, and 3) 0.7μl at AP +0.1, ML +/- 0.8, DV −1.6). AP coordinates were taken from bregma, ML coordinates were taken from the sagittal sinus and DV coordinates were taken from dura.

In Experiment 2, 10 animals received rejections of AAV5-CaMKIIa-hM4Di-mCherry (titer 9.5×10^12 GC/ml; Addgene, MA, USA), while eight animals received injections of a non-DREADD expressing viral control AAV5-CaMKIIa-EGFP (titer 4.3×10^12 GC/ml; Addgene, MA, USA), All animals received three viral injections in the anterior cingulate cortex in each hemisphere as follows (skull at +5.0mm to horizontal plane): 1) 0.35μl at AP +2.1, ML +/-0.8, DV −1.2; 2) 0.65μl at AP +1.4, ML +/- 0.8, DV −1.6, and 3) 0.65μl at AP +0.7, ML +/- 0.8, DV - These injection volumes were as Experiment 1.

To target anterior cingulate efferents to the anterior thalamic nuclei animals, all animals in Experiment 2 were also implanted with guide cannulae aimed at the anterior thalamic nuclei. A craniotomy was drilled in each hemisphere and bilateral guide cannulae (Plastics One, Roanoke, VA, USA) were implanted (26 gauge, cut to a length of 5.4 mm, center to center distance of 2 mm) in the anterior thalamic nuclei at the following coordinates: AP: - 0.1, ML: +/- 1.0, DV: −4.6 (mm from bregma). Cannulae were held in place by bone cement (Zimmer Biomet, Winterthur, Switzerland) and anchored to the skull with four fixing screws (Precision Technology Supplies, East Grinstead, UK). Removable obturators (Plastic One, Roanoke, VA, USA) were inserted into the guide cannulae to prevent the cannulae from blocking.

For all animals (Experiments 1 and 2), the surgical site was closed using sutures and the analgesic bupivacaine (Pfizer, Walton Oaks, UK) was injected between the suture sites. A topical antibiotic powder Clindamycin (Pfizer, Walton Oaks, UK) was then applied to the site. Animals were administered a subcutaneous injection of glucose-saline (5ml) for fluid replacement before being placed in a recovery chamber until they regained consciousness. Animals were monitored carefully postoperatively with food available *ad libitum* until they had fully recovered, with behavioral testing commencing fourteen days post-surgery.

### Behavior

#### Apparatus

All pre-training and testing took place in a black Perspex arena which measured 69.5cm long, 40.5cm wide and 18.6cm tall (Figure 1). One end of the testing arena comprised two individual chambers that were separated from the remaining open space by black Perspex panels that could be removed by the experimenter to allow access. Each of the three compartments had a hinged, transparent Perspex lid. In each of the two smaller compartments was a circular glass pot (75mm diameter, 45mm height) that contained the digging media. Against the opposite wall, in the larger compartment, there was an identical glass pot containing water.

**Figure 1.**
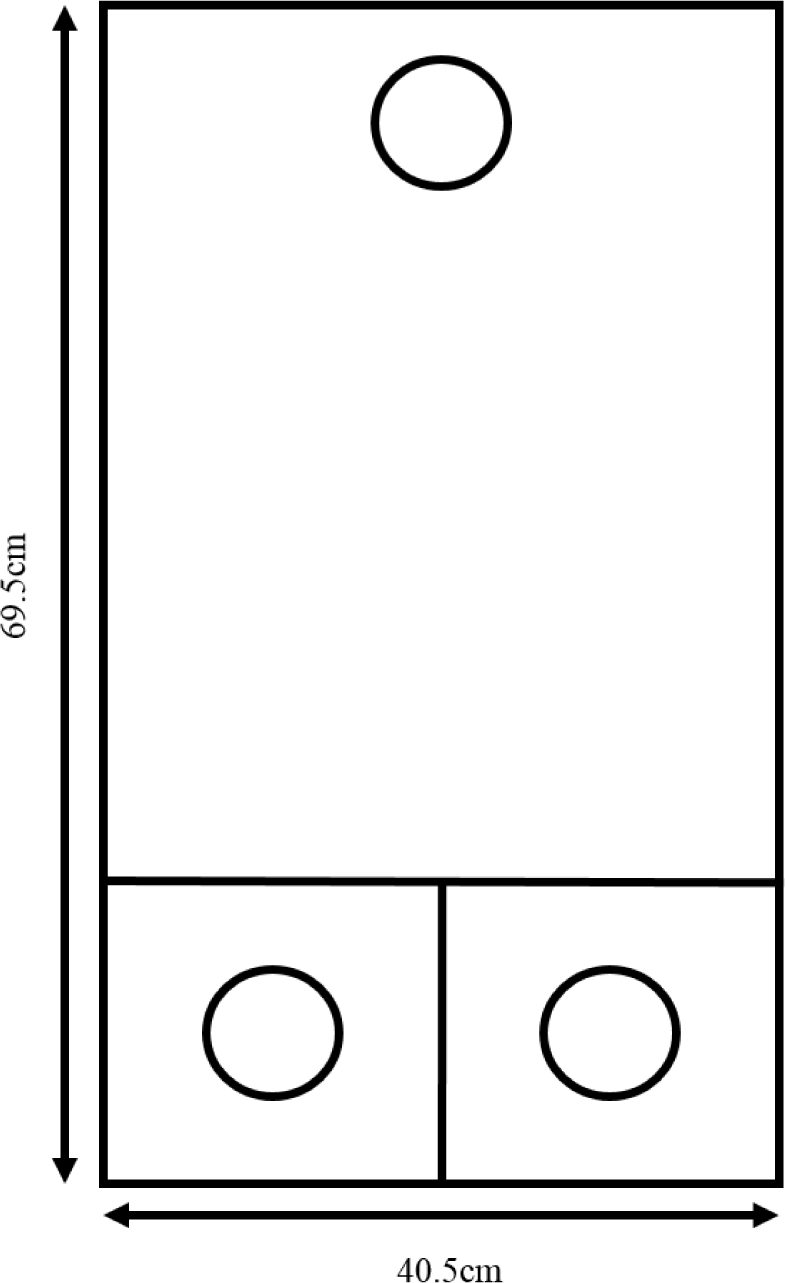
Schematic diagram of the test apparatus used to run the attentional set-shifting task. Approximately one quarter of the space is divided into two smaller chambers, separated from the remaining open space by removable Perspex panels. Each small chamber contains a glass pot containing the digging medium and the remaining, larger open space contains an identical glass pot filled with water.

#### Pre-training

Two weeks after surgery, animals underwent 3 days of pre-training. On the first day of pre-training animals were habituated to the arena for 10 minutes, with access to all three chambers and no glass pots present. On the second day of pre-training, all three glass pots were in place, with the two glass pots in the smaller chambers filled with bedding sawdust. Panels were removed providing access to alternating chambers across trials to prevent the formation of a side bias. On the first trial, half a Cheerio (Nestle, Glasgow, UK) was placed on top of the sawdust and it was progressively buried in subsequent trials to teach animals to dig for the food reward. This ability was typically acquired within 10 trials. The day before testing took place, animals were pre-exposed to the test stimuli (Table 1). Each odor was presented with bedding sawdust and each digging media was presented without odor. Animals retrieved half a buried cheerio from each pot of sawdust laced with odor and each pot of odorless digging media, once in each chamber.

**Table 1.**
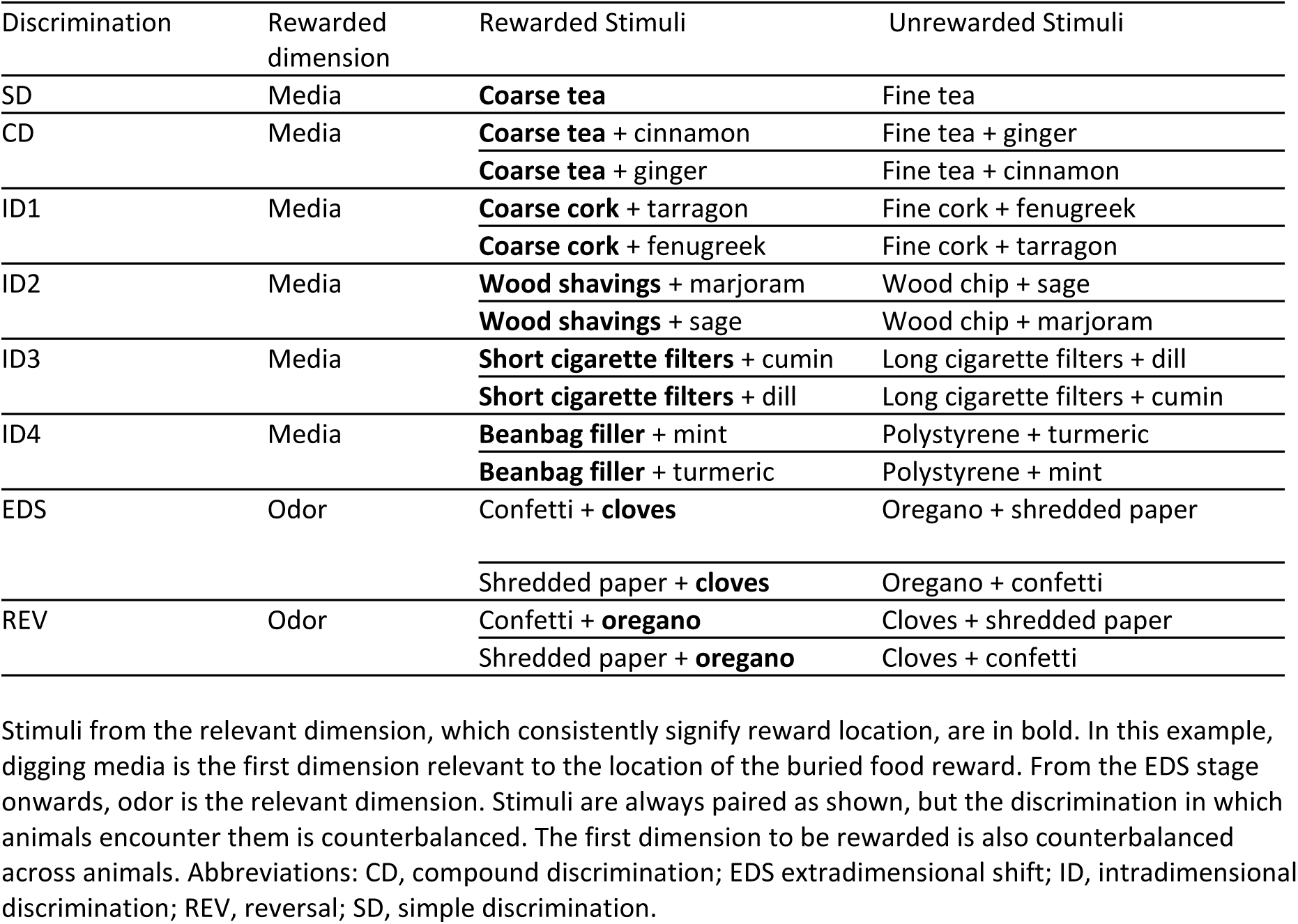
Depiction of a possible order of stimulus pairings in the attentional set-shifting task.

#### DREADD activation

Three weeks post-surgery, behavioral testing on the attentional set-shifting task began. Prior to testing, the DREADDS were activated by clozapine (Gomez et al., 2017; Tan et al., 2020) either by systemic injection (Experiment 1) or intracranial infusion (Experiment 2).

In Experiment 1, clozapine dihydrochloride (Hello Bio, Bristol, UK) was dissolved in sterile saline. Twenty minutes before the test began, all rats received an intraperitoneal (I.P.) injection of clozapine dihydrochloride at a dose of 4 mg/kg (as salt).

In Experiment 2, clozapine dihydrochloride (Hello Bio, Bristol, UK) was dissolved in sterile saline at a dose of 1 µg/µl (as salt). Fifteen minutes prior to testing, rats were lightly restrained, the obturators removed, and 33-gage stainless steel infusion cannulae (Plastic One, Roanoke, VA, USA) that projected 2.0 mm beyond the tip of the guide cannulae were inserted. Each pair of infusion cannula was connected to two 5 μl Hamilton syringes (Bonaduz, Switzerland) mounted on two infusions pumps (Harvard Apparatus Ltd, Kent, UK). A volume of 0.25 μl per hemisphere was infused over 1 minute. The infusion cannulae were left *in situ* for a further 1 minute to allow absorption of the bolus. The infusion cannulae were then removed and the obturators replaced.

#### Behavioral Training: Attentional set-shifting task

Following activation of the DREADDS, the rats received a single test session in the arena (Figure 1). The glass pots in the two smaller compartments of the arena were filled with different stimuli pairs (Table 1). Only one pot contained the buried food reward (half a Cheerio, Nestle, Glasgow, UK). Animals encountered a sequence of discriminations requiring them to learn to select the correct stimulus in order to retrieve the food reward, before beginning the next discrimination.

At the beginning of each trial, the dividing panels were removed allowing the animal access to the two smaller compartments. The compartment of the correct pot was pseudorandomly allocated in each trial. If the animal dug in the correct pot, defined as breaking the surface of the digging media with paws or nose, it could retrieve the reward. For the first four trials of each discrimination, the animal was allowed access to the correct compartment to retrieve the reward following an initial dig in the incorrect pot. Thereafter, if the animal dug in the incorrect pot, access to the correct compartment was blocked. The inter-trial interval lasted approximately five seconds during which time the pots were rebaited. Once the animal had acquired a discrimination, quantified by six consecutive correct digs, it moved on to the next discrimination.

There were eight consecutive discriminations (Table 1). For the initial six discriminations, one dimension, e.g., type of digging media, consistently predicted reinforcement. The session began with a ‘simple discrimination’ (SD) where two distinct digging medias were discriminated. Next, in the ‘compound discrimination’ (CD), the same digging medium continued to be rewarded but stimuli from another dimension (odors) were added to create two different discrimination types (Table 1). For the next four discriminations (ID1-4) the rewarded dimension remained constant (i.e., digging media) but the particular stimuli changed from discrimination to discrimination. Throughout, the other dimension (odor) was present but no individual odor consistently predicted reward. Consequently, improved performance across ID1 to ID4 provides a measure of attentional-set formation, reflecting growing attention to the relevant domain.

For the extradimensional-shift (ED), rats were now required to switch dimensions to solve the discrimination, e.g., from a particular digging medium to a particular odor (Table 1). Finally, rats received a ‘reversal’ in which the particular pair of stimuli in the rewarded dimension had their reward contingencies reversed (Table 1). All testing contingencies were counterbalanced across animals in each group, so that half began with odor as the rewarded dimension.

#### Second extradimensional-shift (spatial)

Approximately two weeks after the first behavioral test, rats completed a second series of discriminations. These took place in the same apparatus and followed the same procedure but comprised four discriminations (Table 2). Prior to this additional test, DREADDS were activated as described above. First, the animals acquired a CD followed by an ID. The rewarded dimension in the first two discriminations (CD, ID) matched that last rewarded in the first behavioral study (i.e. in the ED and Reversal). Next, there was an extradimensional-shift based on the spatial location of the digging pots (EDSpatial, Table 2). Here, for the first time, whether the pot was located in the left or the right chamber became the critical rewarded feature. Both odor and type of digging media were now irrelevant. Following acquisition of the spatial task, the left/right contingencies were reversed (REVspatial, Table 2).

**Table 2.** Possible order of stimulus pairings in the second attentional set-shifting task.

| Discrimination | Rewarded dimension | Rewarded Stimuli | Unrewarded Stimuli |
| --- | --- | --- | --- |
| CD | Odor | **Paprika** + short wire | Coriander + long wire |
|  |  | **Paprika** + long wire | Coriander + short wire |
| ID | Odor | **Lemongrass** + buttons | Nutmeg + beads |
|  |  | **Lemongrass** + beads | Nutmeg + buttons |
| EDSpatial | Spatial location | Lemongrass + buttons (**left chamber**) | Nutmeg + beads (right chamber) |
|  |  | Lemongrass + beads (**left chamber**) | Nutmeg + buttons (right chamber) |
|  |  | Nutmeg + beads (**left chamber**) | Lemongrass + buttons (right chamber) |
|  |  | Nutmeg + buttons (**left chamber**) | Lemongrass + beads (right chamber) |
| REVSpatial | Spatial location | Lemongrass + buttons (**right chamber**) | Nutmeg + beads (left chamber) |
|  |  | Lemongrass + beads (**right chamber**) | Nutmeg + buttons (left chamber) |
|  |  | Nutmeg + beads (**right chamber**) | Lemongrass + buttons (right chamber) |
|  |  | Nutmeg + buttons (**right chamber**) | Lemongrass + beads (right chamber) |

#### Mapping of c-fos expression following DREADD activation (Experiment 1 only)

One week after the completion of behavioral testing, the *in-vivo* influence of DREADDs was investigated by comparing group differences in the expression of the immediate-early gene c-*fos*. This gene is an indirect marker of neuronal activity (Chaudhuri, 1997; Tischmeyer and Grimm, 1999; Guzowski et al., 2005) and is expressed in multiple brain regions, including the anterior cingulate cortex and anterior thalamic nuclei following exposure to novelty (Zhu et al., 1995; Vann et al., 2000; Wirtshafter, 2005).

Rats received an I.P. injection of clozapine dihydrochloride (4mg/kg as salt; HelloBio, Bristol, UK) to activate DREADDS and were placed in a cage in a dark holding room for 20 minutes. Rats were then were placed in two novel environments, each for 15 minutes. The first was a large square open field arena measuring 100cm long, 100cm wide and 45cm tall which was filled with bedding sawdust and six novels objects. The second was a bow-tie maze (Albasser et al., 2010) which measured 120cm long, 50cm wide and 50cm tall and was filled with bedding sawdust and food rewards (Cheerios, Nestle, Glasgow, UK). Rats were then returned to the dark holding room for 90 minutes, to allow for neuronal Fos expression (Bisler et al., 2002) before perfusion.

### Histology

After completion of the behavioral protocols, animals received an I.P. injection of a lethal dose of sodium pentobarbital (2ml/kg, Euthatal, Marial Animal Health, Harlow, Essex, UK) and were transcardially perfused with 0.1M phosphate-buffered saline (PBS), followed by 4% paraformaldehyde in 0.1M PBS (PFA). Brains were removed, post-fixed in PFA for 2 hours and then placed in 25% sucrose solution for 24 hours at room temperature on a stirring plate.

Brains were cut into 40 μm coronal sections using a freezing microtome (8000 sledge microtome, Bright Instruments, Luton, UK) and a series of 1 in 4 sections was collected in PBS for fluorescence analysis (Experiments 1 and 2) as well as Fos analysis (Experiment 1 only). To verify cannulae placements, an additional series was collected for cresyl staining (Experiment 2)

#### DREADDs expression immunohistochemistry

Immunohistochemistry was carried out on the tissue to enhance the fluorescence signal of mCherry (DREADD groups). The first series of sections was transferred from PBS into a blocking solution of 5% normal goat serum (NGS) in Phosphate Buffered Saline with Tritonx-1000 (PBST) and incubated for 1 hour. The sections were then transferred into the primary antibody solution of rabbit-anti-mCherry (Abcam, Cambridge, UK) at a dilution of 1:1000 in PBST with 1% NGS and incubated for 24 hours. Sections were washed four times in PBST and transferred to a secondary antibody solution of goat-anti-rabbit (Dylight Alexa flour 594, Vector Laboratories, Peterborough, UK) or at a dilution of 1:200 at PBST. From this point onwards the sections were kept in the dark. Sections were incubated for 1 hour before being washed three times in PBS. Sections were mounted onto gelatin subbed glass slides and were allowed to dry overnight before being immersed in xylene and coverslipped using DPX (Thermo Fisher Scientific, Loughborough, UK). All incubations were on a stirring plate at room temperature and all washes were for 10 minutes. Virus expression was analyzed using a fluorescent Leica DM5000B microscope with a Leica DFC310 FX camera. Images were collected from the anterior cingulate cortex, selected efferent targets, and a comparison cortical area, secondary somatosensory cortex.

#### Fos Immunohistochemistry (Experiment 1 only)

Sections were washed four times in PBS, once in a peroxidase block (0.3% hydrogen peroxidase in PBST) and four times in PBST. The sections were then transferred to a blocking solution of 3% NGS in PBST and incubated for 1 hour. The sections were then transferred to a primary antibody solution of rabbit-anti-c-*fos* (Millipore, Watford, UK) at a dilution of 1:5000 in PBST and incubated for 10 minutes, followed by 48 hours at 4°C in a refrigerator. Sections were washed four times in PBST and transferred to a secondary antibody solution of goat-anti-rabbit (Vector Laboratories, Peterborough, UK) at a dilution of 1:200 in 1.5% NGS in PBST. Sections were incubated for 2 hours before being washed four times in PBST and transferred to an avidin/biotinylated enzyme complex (Vectastain ABC HRP kit, Vector Laboratories, Peterborough, UK) in PBST for 1 hour. Sections were washed four times in PBST and twice in a Tris buffer (0.6% trisma base in distilled water). Sections were then immersed in a DAB solution (DAB peroxidase HRP substrate kit, Vector Laboratories, Peterborough, UK) for 1-2 minutes before the reaction was stopped with cold PBS. The sections were mounted onto gelatin-subbed glass slides and allowed to dry overnight before being immersed in xylene and coverslipped using DPX. All incubations were on a stirring plate at room temperature and all washes were for 10 minutes.

#### Fos-positive cell counts

Fos-positive cells (diameter 4-20μm, sphericity 0.1-1.0, stained above a grayscale threshold 60 units below peak grey value) were analyzed using a DMRB microscope, an Olympus DP73 camera and cellSens Dimension software (version 1.8.1, Olympus Corporation). Images were collected from consecutive sections (each 120μm apart) using a 5x objective lens. For each hemisphere in each case, eight images were generated for the anterior cingulate cortex and four for both prelimbic cortex and the anterior thalamic nuclei. Three further images were generated for secondary somatosensory cortex, considered a control region as it neither neighbours nor has known interconnectivity with anterior cingulate cortex. For each case, a mean Fos count was generated for images of each region of interest: dorsal anterior cingulate cortex (Cg1), ventral anterior cingulate cortex (Cg2), prelimbic cortex, anteromedial (AM) thalamic nuclei, anteroventral (AV) thalamic nuclei and secondary somatosensory cortex (S2).

### Data Analysis

For the first behavioral task, an initial ANOVA tested for any effects of rewarded dimension (whether rats required to attend to odor or digging media to solve the first discriminations differed), with stage (eight levels) as a within-subjects factor, and first dimension (two levels) and group (two levels) as between-subjects factors. Provided no main effect of rewarded dimension and no interactions between this factor and group were found, data were pooled across dimensions for all subsequent analyses.

A two-way ANOVA examined mean trials to criterion for all eight stages of the attentional set-shifting task, with stage (eight levels) as a within-subjects factor and group (two levels) as a between-subjects factor. ID1-ID4 were examined for intradimensional-set acquisition, while ID4 versus ED assessed shift effectiveness. Following significant interactions, simple effects based on the pooled-error term were used to explore group differences (Howell, 2010). The ‘shift-cost’ was calculated as the difference between the mean trials to criterion for ID1-ID4 and for acquiring the ED (Chase et al., 2012). One-sample t-tests (two tailed) assessed whether the ‘shift cost’ was higher or lower than 0. Errors to criterion were also recorded, giving the same pattern of results across experiments.

For the spatial extradimensional task, an initial ANOVA checked for any effects of rewarded chamber, whether rats were required to dig in the left or right chamber to solve the spatial extradimensional discrimination, on performance. This included stage (four levels) as a within-subjects factor, and first chamber (two levels) and group (two levels) as between-subjects factors. Null results allowed the data to be pooled across dimensions for all subsequent analyses. Next, a two-way ANOVA with stage (four) as a within-subjects factor and group (two levels) as a between-subjects factor examined group differences in mean trials to criterion. Significant interactions were investigated using simple effects based on the pooled-error term. Shift-cost was the difference in the trials to criterion for ID and EDSpatial. One-sample t-tests (two tailed) assessed whether the ‘shift cost’ was higher or lower than 0.

All analyses used JASP computer software (version 0.11.1, Amsterdam, The Netherlands). Data were initially checked for normality using the Shapiro-Wilk test. Mauchly’s test for sphericity was considered and, where violated, Greenhouse-Geuisser corrections were applied to the degrees of freedom. The alpha level was set at p<.05 throughout. To give an estimate of effect size, partial eta squared is reported for all significant main effects and interactions.

A two-way ANOVA examined mean Fos-positive cell counts in the cortical regions of interest, with region (three levels, Cg1, Cg2, PrL) as a within-subjects factor and group (two levels) as a between-subjects factor (Experiment 1 only). A one-way ANOVA was conducted on Fos-positive cell counts in the control cortical region, secondary somatosensory cortex (S2), with group (two levels) as a between-subjects factor. Finally, a two-way ANOVA examined counts in the anterior thalamic nuclei with region (two levels, AM and AV) as a within-subjects factor and group (two levels) as a between-subjects factor. Where interactions were found between region and group, simple effects based on the pooled-error term explored group differences.

## Results

### Experiment 1A: Anterior cingulate cortex intradimensional and extradimensional set-shifting

#### Histology

Two animals, one from each group, were excluded from the analysis due to a lack of expression of the virus in the anterior cingulate cortex. In the remaining animals, analysis confirmed that both viruses were concentrated in the dorsal aspect of the anterior cingulate cortex, Cg1, with some spread into ventral anterior cingulate cortex, Cg2 (Figure 2). Only limited virus reached the edges of the prelimbic or retrosplenial cortices.

**Figure 2.**
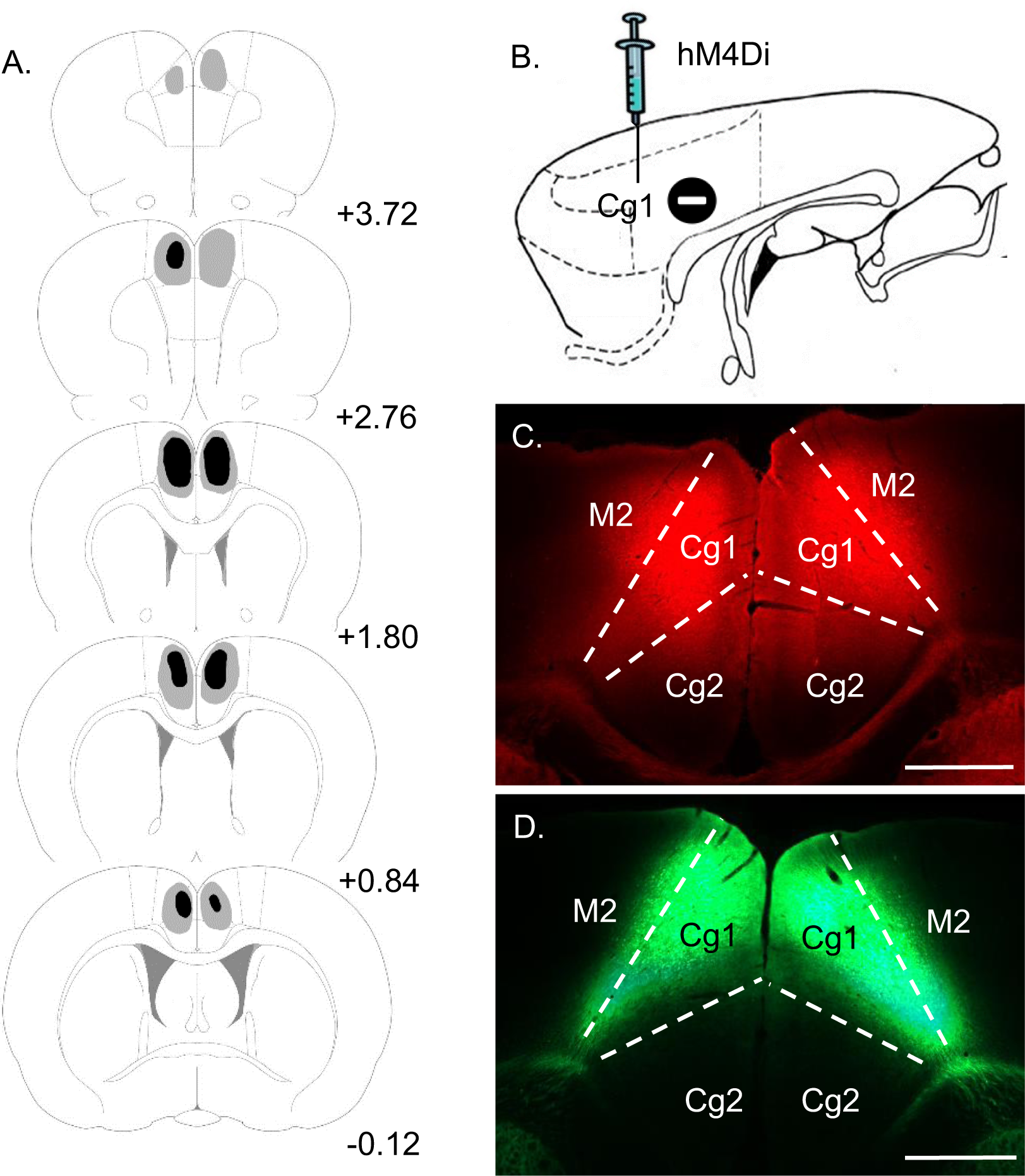
Summary of virus expression in the iDRD (hM4Di) and control groups. **A**. Diagrammatic coronal reconstructions showing the individual cases with the largest (grey) and smallest (black) expression of mCherry in the iDRD group. Numbers refer to the distance (mm) from Bregma (adapted from Paxinos & Watson, 2005). **B**. Sagittal schematic of rat forebrain. **C**. iDRD (hM4Di) mCherry expression in dorsal anterior cingulate cortex (Cg1). **D**. GFP expression in the control group. All animals in both groups displayed robust virus expression centered in Cg1 and its efferent targets. Scale bars 1mm. Other abbreviations: Cg2, ventral anterior cingulate cortex; M2, secondary motor cortex.

#### Behavior

The initial stimulus contingency (odor or media rewarded) did not affect overall performance (F<1), so all data were combined before testing for intradimensional set-formation. For the controls, trials to criterion decreased from ID1 to ID4, the hallmark of ID acquisition (t_(8)_=2.58, p=.033;

Figure 3A). In contrast, the iDRD group failed to improve across these same stages (t_10)_=0.11, p=.91) indicating the lack of an ID set. Further comparisons not only showed that the iDRD group impaired intradimensional set-formation but also facilitated set-shifting (group by eight task stages, interaction F_(7,126)_=3.72, p=.001, η_p_^2^ = .171).

**Figure 3.**
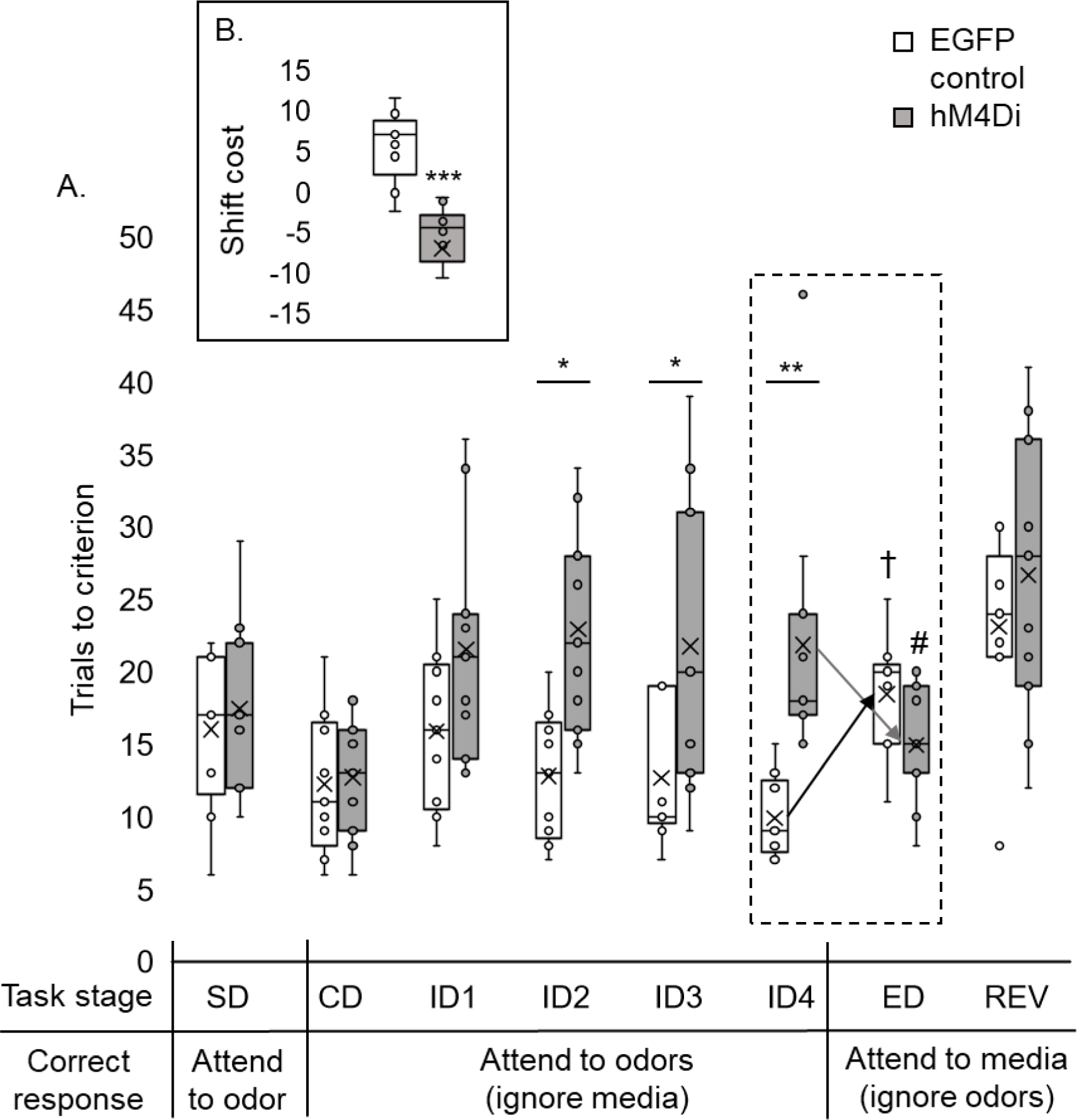
DREADD-mediated inhibition of anterior cingulate cortex efferents (iDRD) impairs intradimensional set-formation but facilitates extradimensional set-shifting. **A**. iDRD (hM4Di) impaired intradimensional set-formation but facilitated the extradimensional-shift (ED). The iDRD group required more trials to solve several individual intradimensional (ID) stages (than the control group, * p<.05, ** p<.01). While the control group required more trials to solve the extradimensional-shift than ID4 († p<.001), the iDRD group took less (than ID4, # p<.05; group interaction, p<.001). **B**. iDRD animals shifted faster than controls (*** p<.001, difference between mean trials to criterion for ID1-4 and ED). While controls had a shift-cost (higher than zero, p<.01), the iDRD group had a shift-benefit (p<.01). The center line in each box represents the median, while the X in each box represents the mean. The box extends between the first and third quartile. Upper and lower whiskers extend 1.5 times the interquartile range. Abbreviations: CD, compound discrimination; REV, reversal; SD, simple discrimination. Note, all discriminations were counterbalanced.

The iDRD group required more trials to solve individual intradimensional stages (ID2 [F_(1,18)_=6.78, p=.018], ID3 [F_(1,18)_=5.42, p=.032] and ID4 [F_(1,18)_=9.40, p=0.007]) than the controls. In contrast, while the controls, as expected, required more trials to solve the extradimensional-shift than ID4 (F_(1,18)_=37.59, p<.001), the iDRD group required fewer trials than for ID4 (F_(1,18)_=7.80, p=.012; interaction (F_(1,18)_= 26.21, p<.001, η_p_^2^ = .593).

Set-shifting (ID to ED) was also examined by considering the ‘shift-cost’, based on the difference in mean trials to criterion across ID1-4 and trials for the ED (Chase et al., 2012; Wright et al., 2015). This analysis revealed a revealed a difference between the two groups (F_(1,18)_= 23.21, p<.001, η_p_^2^ = .563, Figure 3B). While the shift-cost of controls was higher than zero (one-sample t_(8)_=3.75, p=.006), the iDRD group displayed a shift-benefit, i.e., lower than zero (t_(10)_=-3.47, p=.006). Finally, there was no evidence of a group difference on the reversal condition (REV, F<1), nor was there a difference in the mean times taken to complete a trial across the task (F<1).

##### Second extradimensional-shift (spatial)

Rats received a further behavioral test (Wright et al., 2015) to test the generality of the set-shifting facilitation. Now, for the first time, spatial position determined reward (Table 2). There were no group differences on the initial two discriminations (CD, ID), but the iDRD group were again relatively facilitated on the subsequent dimensional switch (EDspatial, Figure 3C). This facilitation is reflected in the interaction from ID to EDspatial (F_(1,18)_=11.18, p=.004, η_p_^2^=.383). The iDRD group shifted faster than the controls (trials to criterion, EDspatial F_(1,18)_=11.78, p=.003, η_p_^2^ = .396). While controls displayed a shift-cost (t_(8)_=2.79, p=.024; ID to EDspatial), the iDRD group displayed neither a cost nor benefit t_(10)_=-1.64, p=.13). There was no difference in mean time taken per trial between the groups (F<1).

### Experiment 1B

In the iDRD group, higher Fos-positive cell counts were observed in cortical area Cg1 (F_(1,18)_=6.88, p=.017,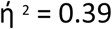), but not Cg2 (F_(1,18)_=2.56, p=.13) or prelimbic cortex (F<1)(group by region interaction [F_(2,36)_=11.16, p<.001, 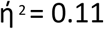])(Figure 5). Further comparisons showed higher Fos-positive cell counts in the iDRD group in both the anteroventral and anteromedial thalamic nuclei (F_(1,18)_=9.73, p=.006, 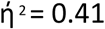). The secondary somatosensory cortex, included as a potential control region, did not show a group difference (F<1).

**Figure 4.**
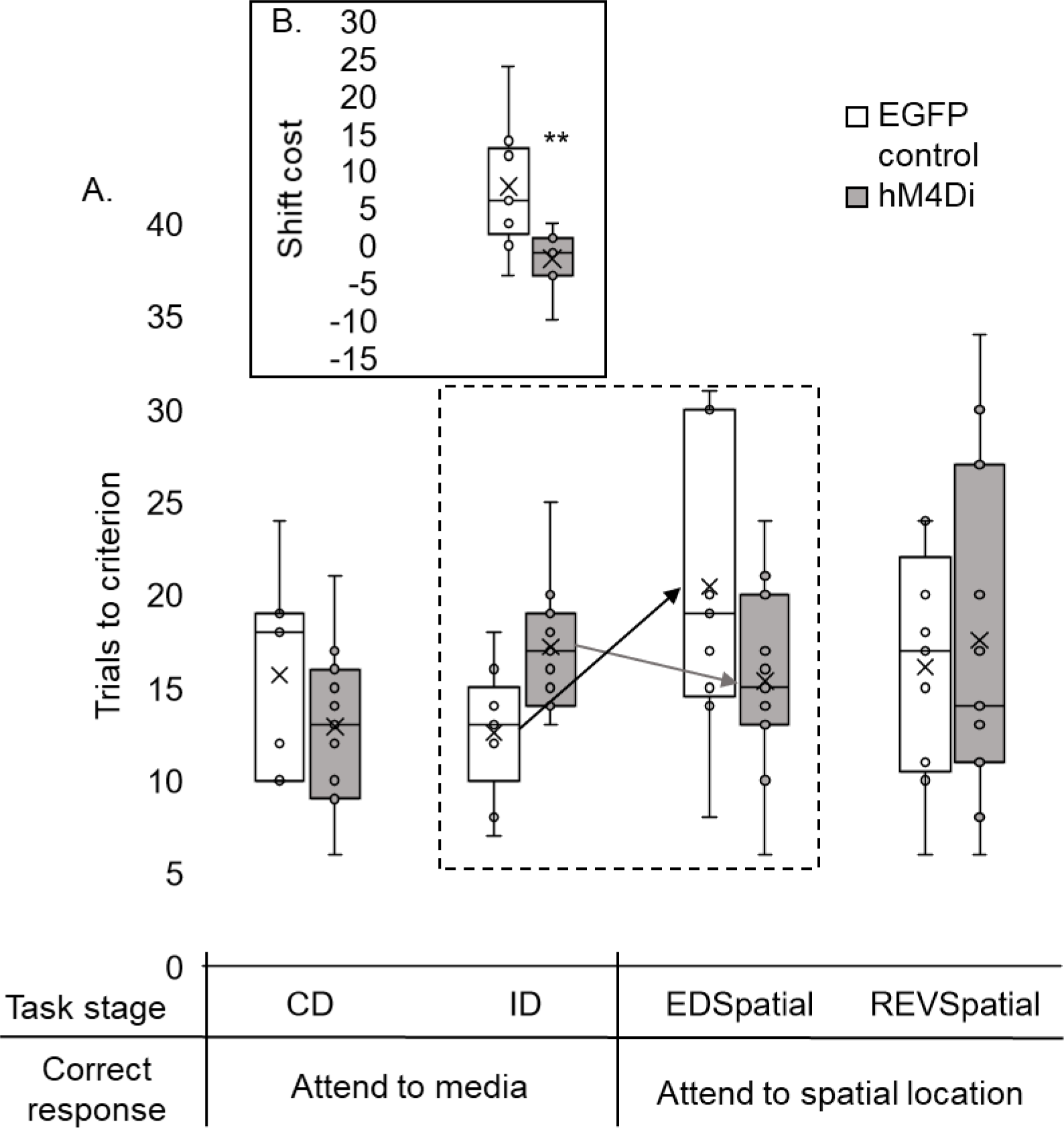
DREADD-mediated inhibition of anterior cingulate cortex efferents (iDRD) facilitates spatial extradimensional set-shifting. A. The control group displayed a shift-cost on an extradimensional-shift based on spatial location (EDSpatial), taking more trials to solve this stage than the intradimensional stage (ID, * p<.05). There was no difference in the number of trials taken to complete these two stages in the iDRD (hM4Di) group. B. The iDRD group shifted faster than controls (** p<.01). The center line in each box represents the median, while the X in each box represents the mean. The box extends between the first and third quartile. Upper and lower whiskers extend 1.5 times the interquartile range. Abbreviations: CD, compound discrimination; REVspatial, spatial reversal. Note, all discriminations were counterbalanced.

**Figure 5.**
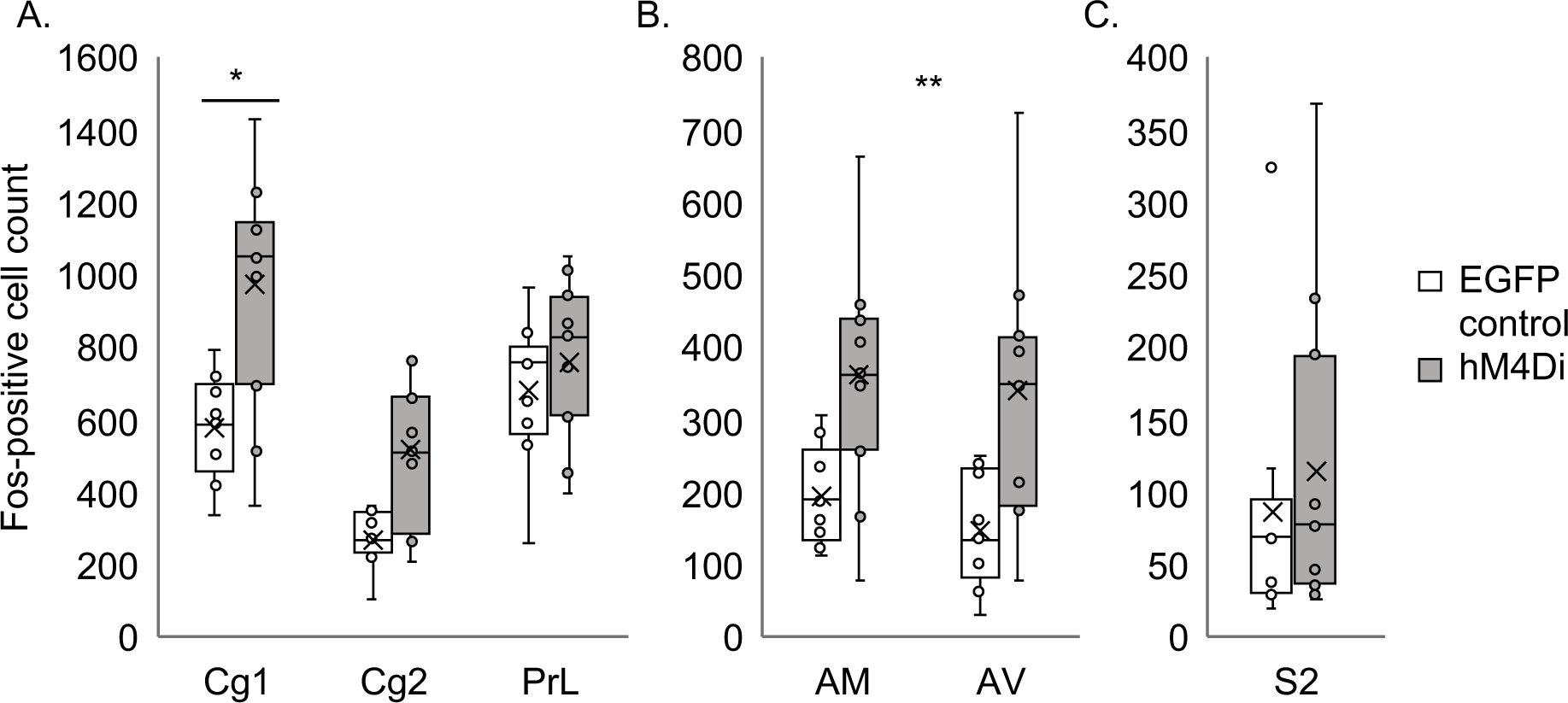
Fos-positive cell counts following DREADD-mediated inhibition of anterior cingulate cortex efferents (iDRD) A. Fos-positive cell counts were higher for the iDRD (hM4Di) group than the control group in Cg1 (*p<.05). There were no differences between the groups in Cg2 or prelimbic cortex (PrL). B. Fos-positive cell counts were higher for the iDRD group than the control group in AM and AV (** p<.01). C. There were no differences between the groups in S2. The center line in each box represents the median, while the X in each box represents the mean. The box extends between the first and third quartile. Upper and lower whiskers extend 1.5 times the interquartile range. Abbreviations: AM, anteromedial thalamic nuclei; AV, anteroventral thalamic nuclei; Cg1, dorsal anterior cingulate cortex; Cg2, ventral anterior cingulate cortex; PrL, prelimbic cortex; S2, somatosensory cortex

### Experiment 2: Anterior cingulate cortex efferents to the anterior thalamic nuclei and intradimensional and extradimensional set-shifting

#### Histology

In both groups, all animals displayed robust virus expression in dorsal anterior cingulate cortex, Cg1, with some spread into ventral anterior cingulate cortex, Cg2 (Figure 6A, C). There was only limited spread to the edges of the prelimbic or retrosplenial cortices. Both viruses showed extensive anterograde transport to sites that receive direct inputs from the anterior cingulate cortex. Among these sites there was dense label in the anteromedial thalamic nucleus and subregions of the anteroventral thalamic nucleus (Figure 6D).

**Figure 6.**
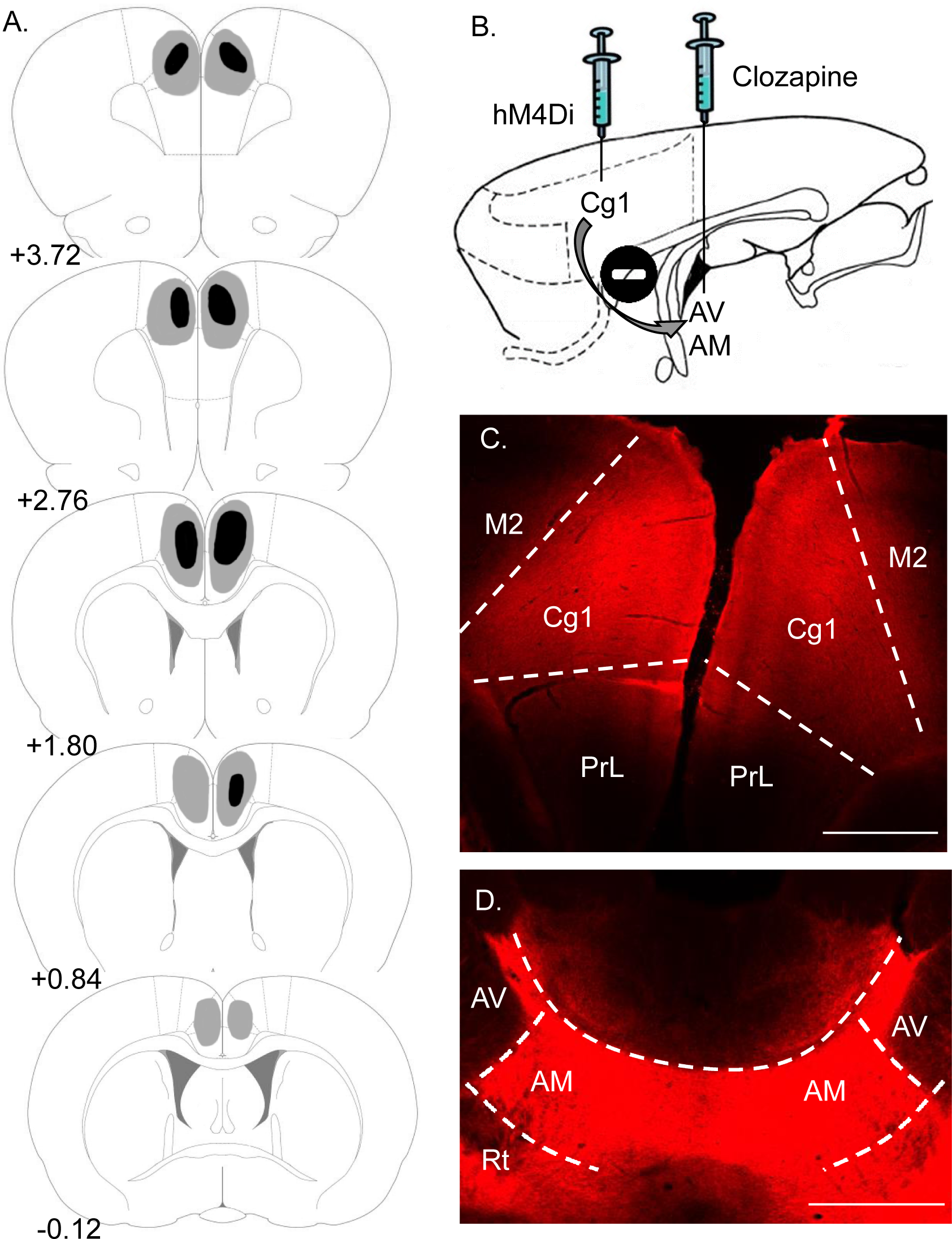
Summary of virus expression in the iDRAccAtn (hM4Di) group. **A**. Diagrammatic coronal reconstructions showing individual cases with largest (grey) and smallest (black) expression of mCherry in the iDRAccAtn group. Numbers refer to the distance (mm) from Bregma (adapted from Paxinos & Watson, 2005). **B**. Sagittal schematic of rat forebrain showing intracerebral clozapine infusion into anteroventral (AV) and anteromedial (AM) thalamic nuclei. **C**. hM4Di mCherry expression in dorsal anterior cingulate cortex (Cg1). **D**. hM4Di mCherry expression in AM and AV. All animals in both groups had robust virus expression in Cg1 and its efferent targets. Scale bars 1mm. Note: GFP expression in control group was comparable to the iDRD group (Figure 2). Other abbreviations: M2, secondary motor cortex; PrL, prelimbic cortex; Rt, reticular thalamic nucleus.

Two animals, one from each group, had cannula tips located outside of the target region of the anterior thalamic nuclei and were excluded from analysis. Figure 7 illustrates the location of the cannulae tips in the remaining animals, identified by histological analysis. The injectors had a 2mm projection, resulting in infusion locations approximately 2mm ventral to the tips of the cannulae.

**Figure 7.**
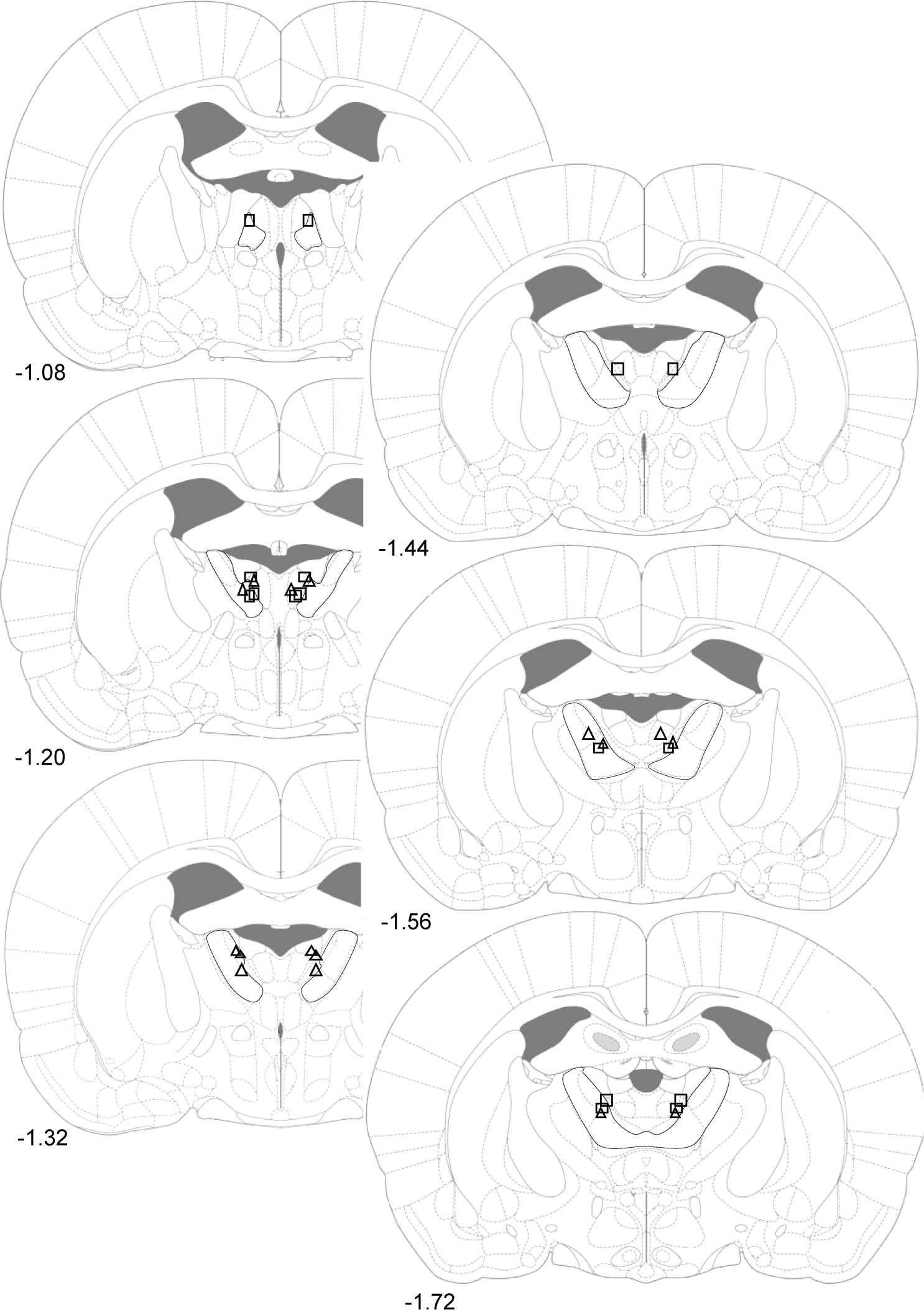
Summary of cannulae placement in the iDRAccAtn and control groups. Diagrammatic reconstructions showing the locations of tips of cannulae aimed at the anterior thalamic nuclei (demarcated with black lines). Triangles represent cases from the iDRAccAtn group and rectangles represent cases from the control group. Numbers refer to the distance (mm) from Bregma (adapted from Paxinos & Watson, 2005).

#### Behavior

The initial stimulus contingency (odor or media rewarded) did not affect overall performance (F_(1,12)_=1.36, p=.27), so all data were combined before testing for intradimensional set-formation. Evidence of set-formation was less clear-cut than in the previous experiment as comparisons between ID1 and ID4 found no significant differences in either controls (t_(6)_=1.80, p=.12) or the iDRAccAtn group (t_(8)_=2.07, p=.07), although ID performance was still disrupted in the iDRAccAtn group (Figure 8). Nevertheless, the controls required more trials to solve the ED than ID4, consistent with prior set-formation (Figure 8A).

**Figure 8.**
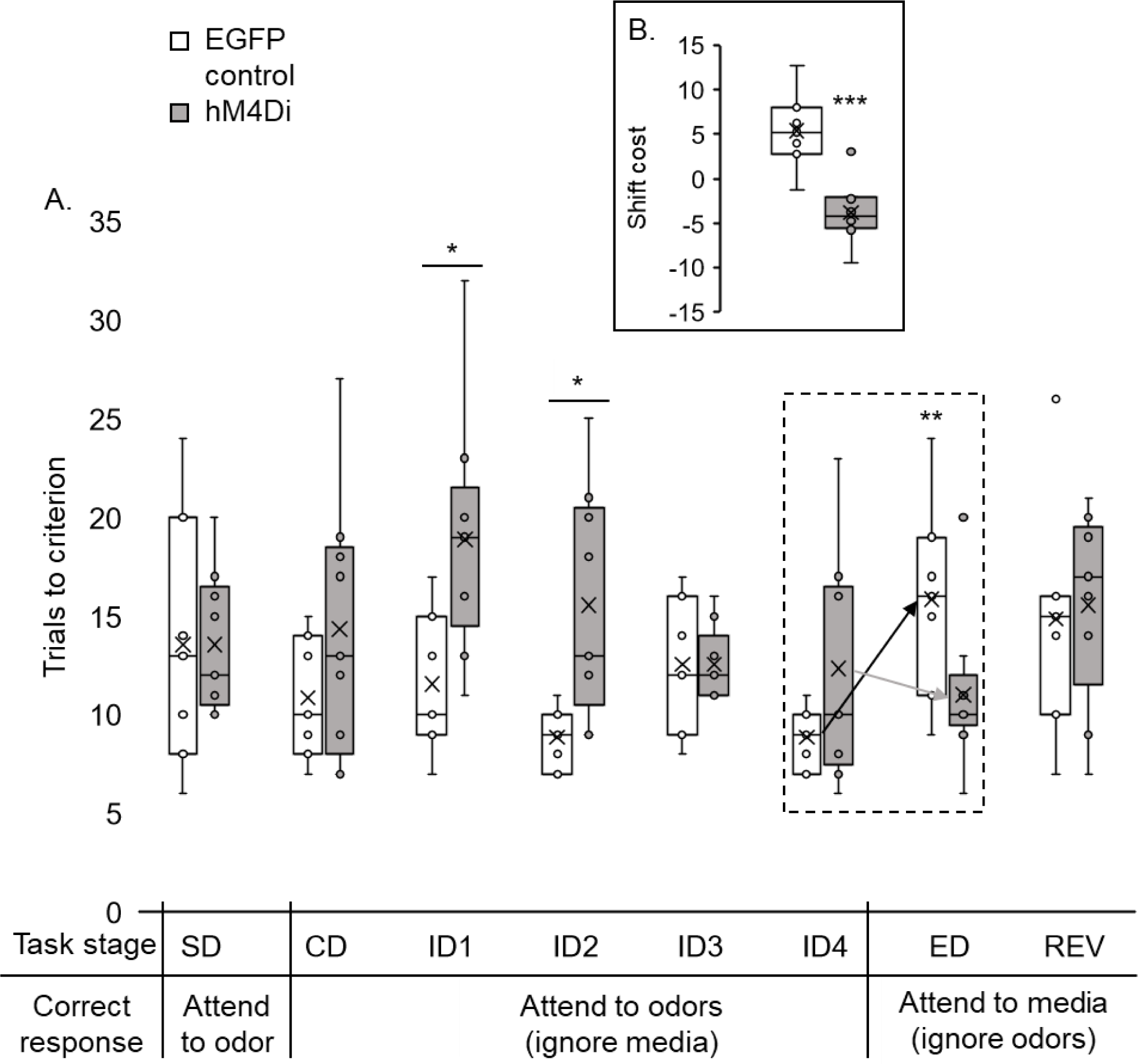
DREADD-mediated inhibition of anterior cingulate cortex efferents to anterior thalamic nuclei (iDRAccAtn) impairs intradimensional set-formation and facilitates extradimensional set-shifting. **A**. iDRAccAtn (hM4Di) animals were impaired at some intradimensional (ID) discriminations (* p<.05) but facilitated the extradimensional-shift (ED). While controls required more trials to solve ED than ID4 (** p<.01) the iDRAccAtn group did not (group interaction, p<.01). B. iDRAccAtn animals shifted faster than controls (*** p<.001, difference between mean trials to criterion for ID1-4 and ED). While controls had a shift-cost (higher than zero, p<.05), the iDRAccAtn group had a shift-benefit (p<.01). The center line in each box represents the median, while the X in each box represents the mean. The box extends between the first and third quartile. Upper and lower whiskers extend 1.5 times the interquartile range. Abbreviations: CD, compound discrimination; REV, reversal; SD, simple discrimination.

Overall comparisons revealed a significant difference in performance between the groups (F_(1,14)_=5.39, p=.036, η_p_^2^ = .278) and an interaction between group and the eight discrimination stages (F_(7,98)_=2.98, p=0.008, η_p_^2^ = .173, Figure 8A). Further analyses showed that iDRAccAtn disrupted learning of ID1 and ID2 (min F_(1,14)_ = 6.87, p=0.02), while extradimensional learning was facilitated in the iDRAccAtn (group interaction ID4, ED, F_(1,14)_= 9.72, p=.009, η_p_^2^ = .41). Controls took more trials to solve the ED (F_(1,14)_ =12.19, p=.004) than the preceding ID4, the iDRAccAtn group showed no difference in the number of trials taken to complete these task stages (F<1). Additional comparisons based on mean trials to criterion for ID1-4 and ED (shift-cost) confirmed these group differences (F_(1,14)_= 22.77, p<.001, η_p_^2^ = .619) (Figure 8B). As expected, controls displayed a shift-cost (t_(6)_ =3.26; p=.017). In contrast, the iDRAccAtn group showed a shift-benefit (t_(8)_=-3.40; p=.009). Finally, performance on the reversal task did not differ (F<1) between groups (Figure 8A).

##### Second extradimensional-shift (spatial)

As in Experiment 1, rats received a second behavioral test including a discrimination where spatial position, for the first time, determined reward (Table 2). There were no group differences on the initial two discriminations (CD, ID), but the iDRAccAtn group were again relatively facilitated on the subsequent dimensional switch (EDspatial, Figure 9). This facilitation is reflected in the interaction from ID to EDspatial (F_(1,14)_=9.91, p=.007, η_p_^2^ = .415). The iDRAccAtn group shifted faster than the controls (trials to criterion, Edspatial F_(1,14)_=15.57, p=.001, η_p_^2^ = .527, Figure 9B). While controls displayed a shift-cost (t_(6)_=5.61, p=.001; ID to EDspatial), the iDRAccAtn group displayed neither a cost nor benefit (t<1). There was no difference in mean time taken per trial between the groups (F<1).

**Figure 9.**
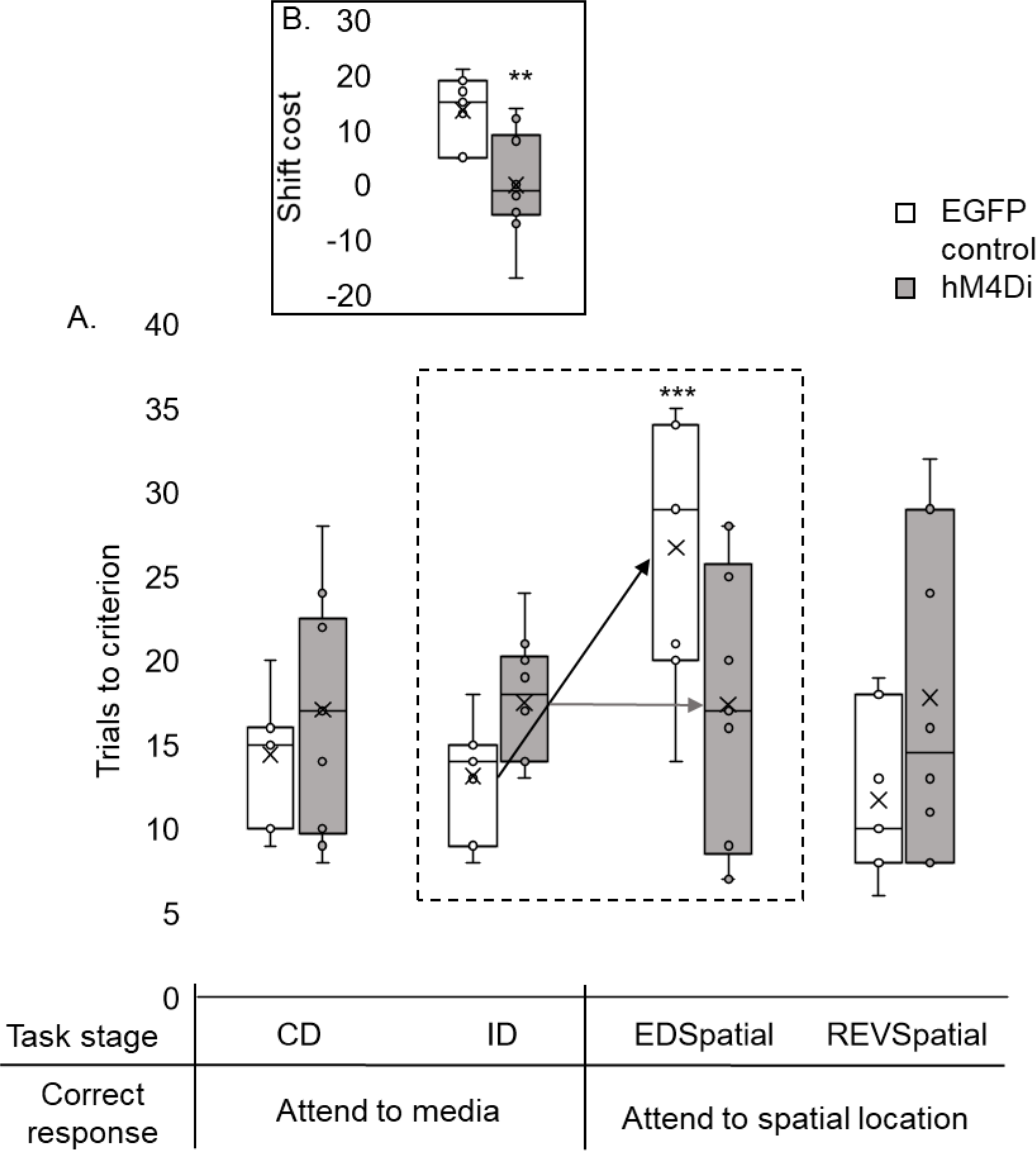
DREADD-mediated inhibition of anterior cingulate cortex efferents to anterior thalamic nuclei (iDRAccAtn) facilitates spatial extradimensional set-shifting. A. The control group displayed a shift-cost on the spatial extradimensional-shift (EDSpatial), taking more trials to solve this discrimination than the intradimensional (ID) stage (*** p<.001). In the iDRAccAtn (hM4Di) group, there was no difference in the number of trials taken to complete these two stages. B. The iDRAccAtn group shifted faster than controls (** p<.01). The center line in each box represents the median, while the X in each box represents the mean. The box extends between the first and third quartile. Upper and lower whiskers extend 1.5 times the interquartile range. Abbreviations: CD, compound discrimination; REVspatial, spatial reversal. Note, all discriminations were counterbalanced.

## Discussion

The present study began by demonstrating that the anterior cingulate cortex (ACC) is required for attentional set-formation in rats. The inhibitory ACC DREADD group failed to show accelerated learning over successive intradimensional-shifts (ID) and was impaired relative to control animals on several of these discriminations. However, when first required to solve a discrimination involving the previously irrelevant stimulus dimension (extradimensional-shift; ED), the same animals showed a significant shift-benefit, requiring fewer trials to solve the ED relative to the proceeding ID stages. This profile of impaired attentional set-formation but facilitated switching reproduces precisely the pattern of performance on the same task by rats with lesions in the anterior thalamic nuclei (ATN) (Wright et al., 2015). Experiment 2 then showed how interactions between the ACC and ATN are required for successful attentional set-formation, as chemogenetic silencing of ACC efferents to the ATN also impaired ID but again facilitated ED shifts.

### Methodological Considerations

A primary consideration relates to whether the administration of clozapine in the EGFP control animals had off-target or non-specific effects that could confound interpretation of the behavioral results (Roth, 2016; Smith et al., 2016). This issue is particularly germane to Experiment 1 which involved systemic injections of the ligand. Although the possibility of off-target effects of the ligand cannot entirely be discounted, there is little evidence to suggest that such effects confound interpretation of the key behavioral findings. For example, comparing the performance of the EGFP groups across Experiments 1 and 2, there was no difference in the magnitude of the key shift-cost measure. Furthermore, the profile of task performance of the EGFP groups in the current experiments closely match those of controls groups previously reported by both our and other groups (Chase et al., 2012; Lindgren et al., 2013; Wright et al., 2015; Powell et al., 2017; Tait et al., 2018). The unique learning profile of the experimental groups (impaired ID but facilitated ED shift performance) is similarly difficult to reconcile with an account framed in terms of off-target effects of the ligand.

Experiment 1B assessed the effects of DREADD activation within the dorsal ACC and key efferent targets on levels of Fos protein. This analysis showed changes, relative to EGFP controls, in dorsal ACC, and in the anteromedial and anteroventral thalamic nuclei. Perhaps unexpectedly, activation by clozapine increased, rather than decreased, Fos levels in these sites. These changes were, however, selective as no Fos differences were observed in ventral ACC, prelimbic or sensorimotor cortex. While studies assessing the effects of inhibitory DREADD activation on network activity have generally reported decreases in c-*fos* expression (Koike et al., 2016; Choi and McNally, 2017), others have found increased activity, as seen in the current experiment (López et al., 2016). Anterior cingulate excitatory neurons innervate the thalamic reticular nucleus, which, in turn, sends inhibitory projections to the anterior thalamic nuclei (Lozsádi, 1994; Gonzalo-Ruiz and Lieberman, 1995; Zikopoulos and Barbas, 2006). DREADD-mediated inhibition of reticular neurons could, therefore, disinhibit anterior thalamic - cortical circuitry. Of particular relevance is how the Fos counts provide independent evidence for abnormal activity associated with DREADD action, activity that was focused in those sites examined in Experiment 2.

### Anterior cingulate cortex, the anterior thalamic nuclei and attentional set-formation

While it has previously been reported that lesions in the orbitofrontal cortex and dorsomedial striatum impair attentional set-formation (Chase et al., 2012; Lindgren et al., 2013), the deficit in attentional set-formation combined with the facilitation of ED shifts seen here after inhibition of the ACC (Experiment 1) and its efferents to the ATN (Experiment 2) is highly unusual. This learning profile is consistent with an initial bias to process poor predictors of reward. Such a learning bias would retard ID shifts when animals are required to preferentially attend to one stimulus dimension that is consistently predictive of reward (e.g. odor) while ignoring other irrelevant stimulus dimensions (e.g. media).

This interpretation, an attentional bias towards poor predictors of reward, was strongly supported by the facilitation of learning seen in the DREADD groups at the ED stage when the hitherto irrelevant dimension now became critical to solving the discrimination. Moreover, the experiments also highlight the generality and robustness of these attentional effects as a further set-shifting facilitation was found for another stimulus class, spatial position. Meanwhile, the selectivity of the current DREADDs effects was underlined by repeated evidence of intact reversal learning (Experiments 1 and 2), where reward contingencies switch but remain within the same stimulus pair, so modality stays neutral. Reversal deficits are, however, seen after orbitofrontal cortex lesions (Dias et al., 1996a, 1996b; Chase et al., 2012) (but see Murray and Rudebeck, 2018). These dissociable profiles highlight how ‘cognitive flexibility’ reflects multiple processes (Kehagia et al., 2010).

Experiment 2 targeted anterior cingulate cortex projections to the anterior thalamic nuclei. The more selective nature of this manipulation helps to explain the less disruptive effects on ID1-4 than those seen in Experiment 1, in which multiple anterior cingulate efferents were compromised. Nevertheless, a significant ‘shift-benefit’ was again observed. These results advance prior evidence that the ATN have an important role in set-formation (Wright et al., 2015), evidence that includes neuropsychological findings (de Bourbon-Teles et al., 2014). This attention role is, however, specific as ATN lesions spare vigilance tasks (Chudasama et al., 2001), despite such tasks being disrupted by ACC lesions (Muir et al., 1996). In turn, these ACC lesion attentional effects are qualitatively different from those associated with the prelimbic and infralimbic cortices (Muir et al., 1996; Koike et al., 2016; Fisher et al., 2019).

The demonstration here that the ACC and its efferents to the ATN mediate attentional set-formation accords with evidence demonstrating a role for the ACC in incorporating reward history with current contingencies to determine action selection (Behrens et al., 2007; Rushworth et al., 2007; Shenhav et al., 2016). For example, lesions in the primate ACC impair the ability to use the past history of actions and their outcome to select appropriate behavioral responses, while neurons in the primate ACC respond during the generation of exploratory actions and the monitoring of outcomes of these actions (Hayden and Platt, 2010). If one function of the ACC is to mediate the relationship between recent actions-outcomes and current behavioral choices (Rushworth et al., 2004, 2007; Quilodran et al., 2008), inhibition of the ACC would be predicted to impair animals’ ability to establish those stimulus-dimensions consistently associated with reward, and with no-reward, respectively. This description matches the pattern of behavior seen in the current experiments. A failure to acquire an attentional-set would be expected to abolish the shift-cost normally seen at the ED stage (Durlach and Mackintosh, 1986). Inhibition of the ACC and its efferents to the ATN did not just abolish the shift-cost associated with the ED, it facilitated switching. The implication, therefore, is that the ACC mediates the relationship between previous actions that are reliably associated with outcomes and current behavioral choices. A further implication is that those systems that modulate attention and responding to inconsistently reinforced action-outcome associations must reside outside the ACC and ATN.

### Conclusions

This facilitation of set-shifting after ACC disruption is the converse of that seen after medial prefrontal cortex lesions. While lesions of medial frontal areas in monkeys and rats spare set-formation, ED switching is protracted, requiring more trials than controls (Dias et al., 1996b, 1996a; Birrell and Brown, 2000). Likewise, patients with frontal lobe damage have difficulty in shifting to a previously irrelevant dimension (Owen et al., 1991). The resulting double dissociations (ACC versus medial prefrontal) on set-formation and set-switching can best be explained by two competing processes (Mackintosh, 1975; Pearce and Hall, 1980; Pearce & Mackintosh, 2010), each with distinct neural underpinnings. While ACC activity would normally promote adherence to a previously successful stimulus class (lost after ACC inhibition), medial prefrontal cortex promotes attentional switching (Sharpe and Killcross, 2014).

These strategies match those separately predicted by learning theorists (Mackintosh, 1975; Pearce and Hall, 1980; Pearce and Mackintosh, 2010). Together, they determine how past learning guides present choice behavior (Pearce and Mackintosh, 2010). It is presumed that the normal activity of the rat ACC, in cooperation with the ATN, is to focus learning and attention on successful reward outcomes while updating internal models of the environment, e.g., intradimensional set-formation. Disruption of this function leads to an initial excessive attention to irrelevant cues which, paradoxically, can facilitate learning when contingencies change. The interplay between these ‘switch’ and ‘stay’ processes closely relates to cognitive inflexibility, a prominent feature of multiple psychiatric conditions (Gold et al., 1997; Merriam et al., 1999; Tsuchiya et al., 2005; Meiran et al., 2011; Polak et al., 2012). The specific difficulty of selecting appropriate stimuli to guide choice behavior has obvious similarities with the symptoms of psychiatric conditions, including depression, schizophrenia, and autism spectrum disorders (Gold et al., 1997; Merriam et al., 1999; Tsuchiya et al., 2005; Meiran et al., 2011), all conditions that display ACC dysfunction (Yücel et al., 2003). One such example concerns how attention to irrelevant cues is related to positive symptoms in schizophrenia (Morris et al., 2013).

## Acknowledgments

We thank E. Amin for her technical support. Funding: This research was supported by a joint grant to JPA and SMOM from the Wellcome Trust (103722/Z14/Z) as well as PhD studentship to EB from the School of Psychology, Cardiff University.

## Author contributions

E.B., A.N., S.OM. & J.A. designed the research; E.B performed the research; E.B. analyzed the data; A.N. & J.A. wrote paper.

